# Developing a behavioral measure of social memory from narrative video

**DOI:** 10.64898/2026.09.09.750409

**Authors:** Taylor Marcus, Didi Ross, Roma Kolluru, Ben Deen

## Abstract

Understanding other people relies on long-term memory: we use learned information about others’ personalities, mental states, and personal histories to make inferences about the causes of their behavior. However, little is known about individual differences in memory for high-level social information. Here, we develop and psychometrically evaluate a behavioral measure of social memory from narrative video content. Participants (*N* = 103) watched the pilot episode of the TV show *Friday Night Lights*, and answered multiple choice questions about characters’ actions, statements, inferred mental states, relationships, and personalities immediately after viewing and following a 21-day delay. The social memory task showed moderate split-half reliability, demonstrating feasibility as an individual difference measure. Memory for information about specific events in the narrative showed a decline across the delay, while memory for abstract information remained stable. Social memory performance was correlated with separate measures of face-name association and verbal episodic memory, but not with face image recognition memory. These results provide a proof of concept for a novel behavioral measure of social memory and suggest that the measure relates to episodic and relational memory but not image recognition.

## Introduction

Human social cognition relies heavily on long-term memory. To make inferences about why people are doing the things they’re doing, and what they might do in the future, we use information stored in memory about others’ personality traits (Fiske et al., 2007; Hastie & Kumar, 1979; Srull & Wyer, 1989), mental states (Ciaramelli et al., 2013; Jara-Ettinger et al., 2016), and relationships (Gershman & Cikara, 2020; Jolly et al., 2023). Likewise, declarative memories are structured by social content. Cognitive models of events are scaffolded by social information: which individuals were present, how they interacted, and how their actions were generated by underlying causes (Zwaan & Radvansky, 1998).

Neuroscientific evidence also suggests a close relationship between systems supporting social cognition and declarative memory. Tasks involving both social cognition (Contreras et al., 2012; Saxe & Kanwisher, 2003) and semantic and episodic memory retrieval (Addis et al., 2007; Binder & Desai, 2011; Szpunar et al., 2007) elicit responses in the default network (DN), a distributed set of brain regions with anatomical and functional connections to the medial temporal lobe (Binder et al., 2009; Vincent et al., 2006). The DN represents diverse social properties, including personality traits (Hassabis et al., 2014; Tamir et al., 2016), mental states (Koster-Hale et al., 2014; Skerry & Saxe, 2015), and relationships (Park et al., 2021; Peer et al., 2021). This system is also sensitive to event structure in social narratives, responding more strongly and reliably to sequences of sentences that build a coherent narrative, compared to unrelated sentences (Ferstl & Von Cramon, 2001; Yarkoni et al., 2008). These properties make the DN well-placed to represent social structure in long-term memory.

Early work argued that overlapping responses to social cognition and memory tasks in the DN reflected the engagement of a single, common system, and a domain-general cognitive operation such as episodic projection (Buckner & Carroll, 2007; Spreng et al., 2009). Recent studies using precision fMRI to capture fine-grained organization in individual brains supports a more nuanced view, showing that the DN comprises at least two parallel, interdigitated networks termed DN-A and B (Braga & Buckner, 2017; DiNicola et al., 2020). DN-A and B show strongly dissociable content domain preferences: DN-A responds selectively to information about places, while DN-B responds selectively to information about people (Deen & Freiwald, 2025). Both networks are sensitive to mnemonic content, with DN-B showing a preference for familiar over unfamiliar people (Deen & Freiwald, 2025). DN-B may therefore constitute a domain-specific component of the corticohippocampal long-term memory system, specialized for social memory.

Evidence for a social memory system in the brain raises the need for a behavioral measure of individual differences in long-term memory for high-level social information. Such a measure could be used to assess the domain-specificity of declarative memory systems at the behavioral level, to relate individual differences in social memory to measures of DN-B function and anatomy, and to measure changes in social memory ability in aging and disease. While memory for personality traits and actions has been extensively studied (Leshikar, 2024; Srull & Wyer, 1989) and some studies have evaluated individual differences (Lassiter et al., 1991; Srull et al., 1985), prior work has not developed a validated measure intended for broader use. Standard measures of long-term memory involve simple, well-controlled stimulus materials such as words or images, lacking social content. In the Rey Auditory Verbal Learning Task (RAVLT), one component of the NIH Toolbox Cognition Battery (Gershon et al., 2013; Weintraub et al., 2013), participants hear a list of words and are asked to repeat them after a delay, measuring verbal episodic memory (Rey, 1964). Other tasks capture memory for basic social information, such as face–name associations (Amariglio et al., 2012; Rentz et al., 2011). However, face-name associations are just one component of the rich social content we learn from experience and use to understand other people.

How can we study individual differences in memory for high-level social information? One promising approach is to assess memory for social content in narrative videos such as television shows and movies. Like our real-world experience, narrative videos present a contextualized and dynamic sequence of complex events (Furman et al., 2007). Narratives in television shows and movies typically center around social content – characters’ actions, interactions, and converesations – and provide rich cues to underlying mental states and personality traits (Levin et al., 2013). Despite their complexity relative to controlled experimental stimuli, narrative videos have been used effectively in research on long-term memory (Delarazan et al., 2023; Furman et al., 2007), social cognition (Krendl et al., 2022), and emotion (Cheong et al., 2023).

The current study develops and validates a behavioral measure of long-term social memory from narrative video content, with the goal of better capturing the demands placed on memory systems in everyday social life. Participants watched the pilot episode of the television show *Friday Night Lights*. Immediately and following a 21-day delay, they completed a social memory task (SMT) comprised of multiple choice questions probing memory for high-level social information: characters’ actions, statements, personality traits, mental states, and relationships. To determine how performance on this social memory task relates to established individual difference measures of long-term memory, we administered the RAVLT and tests for face-name association and image recognition memory. The results demonstrate that the social memory task captures reliable individual differences in memory performance. These differences related to existing laboratory-based measures of episodic memory, but were not fully explained by these measures, supporting the task’s utility as a complementary measure of naturalistic social memory.

## Methods

### Participants

*N* = 103 adults from Tulane University and the greater New Orleans area participated in the study (age 18-22 years old, M_Age_ = 18.93, SD = 1.21, 71 females, 32 males). Per self-report, 76.70% of participants were white, 5.83% were multiracial, 10.68% were Asian, .97% were Hispanic, and 5.83% were Black. Participants were excluded if they had any history of diagnosed psychiatric or neurological condition, uncorrected visual impairment, or if they had previously watched any episodes of *Friday Night Lights*. An additional 53 participants were enrolled but excluded due to non-compliance (e.g., missing the second study visit; *N* = 24), experimenter error (*N* = 4), or technical error (e.g., experiment crashing; *N* = 25). All study procedures were approved by the Tulane University Institutional Review Board, and participants provided written, informed consent.

### Materials and Procedure

Participants completed two in-person testing sessions (Figure 1). At visit 1, participants watched the pilot episode of the show *Friday Night Lights* and completed the first half of the SMT immediately after watching the episode (Figure 2). *Friday Night Lights* was selected because the pilot episode contains rich social and emotional content, a compelling narrative, well-developed characters, and clearly conveyed relationships, and has been effectively used in prior work (Chang et al., 2021; Cheong et al., 2023). After watching the pilot video, participants were asked to rate their subjective enjoyment of the show on a scale of 1-10.

**Figure 1:**
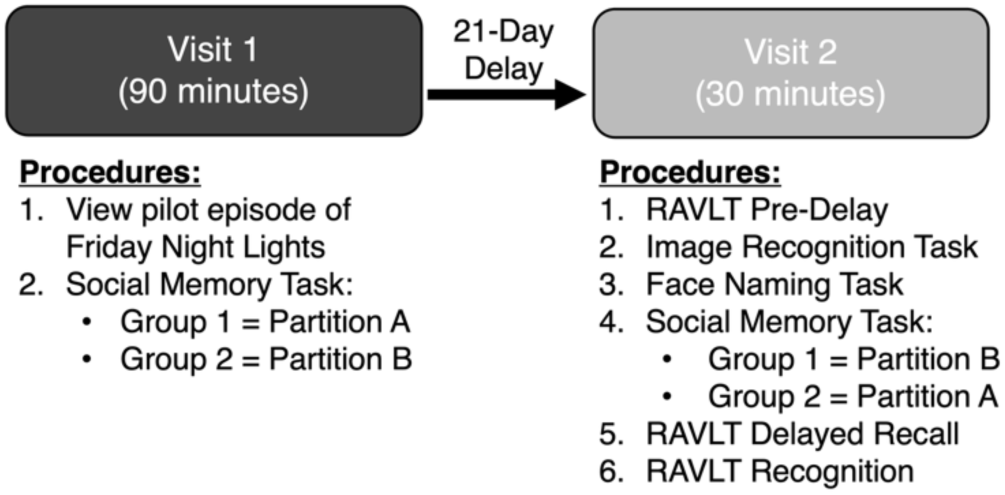
Visit structure.

**Figure 2:**
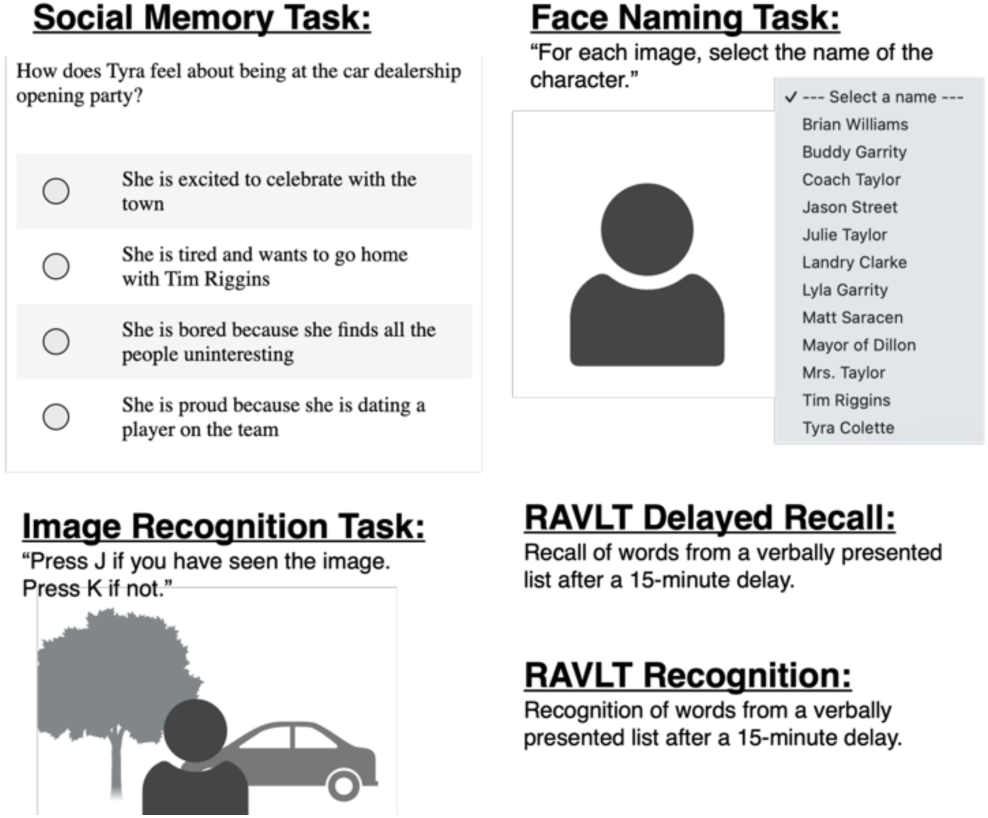
Task descriptions.

Visit 2 was completed three weeks after visit 1 (M_delay_= 21 days, SD = 0) to investigate memory decay. Participants were instructed not to view any additional episodes of the show between visits 1 and 2. At visit 2, participants completed the second half of the SMT along with a battery of memory tasks: the Rey Auditory Verbal Learning Test (RAVLT), an image recognition task, and a face naming task (Figure 2).

### Social memory task

The SMT included 50 four-option multiple-choice questions designed to assess memory for diverse social information. Questions were separated into two types: event-based and abstract. Event-based questions referred to specific events in the narrative, and were further separated into three subtypes: questions about characters’ mental states (e.g. “How does Tyra feel about being at the car dealership opening party?”), actions (e.g. “After seeing Lyla in the hallway of the hospital, what does Julie do?”), and statements (e.g. “Before leaving the house, what does Lyla say to her father?”). Abstract questions referred to generic information not tied to specific events, and were further separated into three subtypes: questions about characters’ mental states (e.g. “What does Lyla think about Tyra?”), personality traits (e.g. “How would you describe Lyla?”), and relationships (e.g. “How does Landry know Julie?”). Note that while the distinction between event-related and abstract information is conceptually related to the distinction between episodic and semantic memory, we use the former as descriptive term for question content without making assumptions about cognitive processes used to answer the questions.

Questions about mental states and personality traits probed participants’ inferences about latent properties that were not directly conveyed by the stimulus, and for which no ground truth answer exists. For these questions, the correct answer was determined by researchers’ judgment. Questions were only included when two researchers (D.R. and B.D.) agreed on the correct answer. As described below (Results), performance on the task with no delay was high across subcategories, demonstrating agreement between researchers and participants.

To probe social memory with and without a delay, the questions were split into two equal partitions (A and B) of 25 questions with balanced question counts by subtype. Participants completed one partition at visit 1 and the other at visit 2. Partition order was counterbalanced across participants to ensure that effects of delay were not driven by question content. Group 1 participants received partition A at visit 1 and B at visit 2, while group 2 participants received partition B at visit 1 and A at visit 2. Group was randomly assigned across participants. Within each visit, question order was randomized for each participant. While conducting the social memory task, participants were given printed document containing images of each character’s face paired with their name. This ensured that participants could link names used in the questions to characters in the show, and was included after pilot testing revealed that participants found the task too difficult without this information.

### Face naming task

In the face naming task, participants were shown face images of characters from the episode they had watched, with twelve characters included. Participants were asked to identify each character by name via a drop-down menu to assess memory for face-name associations. Image order was randomized for each participant. The face naming task was performed prior to the social memory task, ensuring that participants had not yet seen the document linking faces to names used for the SMT.

### Image recognition task

In the image recognition task, participants were shown still images from the show, including both familiar images from the pilot episode and unfamiliar control images from scenes in subsequent episodes that were not seen. Each image contained a single character, and each control image was matched to a familiar image based on the character present. 48 trials were presented in random order. Each trial consisted of image presentation (1.5 seconds) followed by a histogram-matched pink noise visual mask (0.1 seconds) and fixation cross (jittered intertrial interval, 2-3 seconds). Participants responded with a button press during the image, mask, or fixation to indicate whether the image was familiar or unfamiliar. The image recognition task was performed prior to the face naming task.

### Rey Auditory Verbal Learning Test (RAVLT)

The Rey Auditory Verbal Learning Test (RAVLT; Rey, 1964) was used to assess verbal episodic memory. In the acquisition phase, participants were read a list of 15 words repeated over 5 trials and were asked to recall as many words as they could after each trial. On trial 6, participants were read a second, novel list of words and asked to recall as many words as they could. On trial 7, participants were not read a list, but instead asked to reacall as many of the words from the initial list (trials 1-5) as they could. Then, after a 15-20-minute delay, participants were asked to recall the initial list again in a delayed recall trial. During the delay, participants performed the image recognition, face naming, and social memory tasks.

Following the recall trials, the RAVLT also included a recognition task. Participants were given a sheet of words that included both words from the repeated list and distractor words. Participants were asked to identify the words they recognized from the list. Preliminary analyses found that delayed recall and recognition were strongly correlated (*r* = .78). Subsequent analyses therefore focused on recall.

## Results

### Task Performance

Performance on each task was evaluated by comparing accuracy against chance using a *t*- test. Participants performed significantly and substantially above chance on the SMT at both visits. At visit 1, mean accuracy was 90.1% (*SD* = 8.1%, chance 25%, *t*(102) = 81.29, *p* < .001). At visit 2, mean accuracy was 78.1% (*SD* = 10.6%, *t*(102) = 50.89, *p* < .001). Performance patterns were consistent across groups at both visits (Supplementary Results). Face naming task accuracy was significantly above chance (*M* = 68.2%, *SD* = 18.8%, chance 8.3%, *t*(102) = 32.31, *p* < .001), as was image recognition (*M* = 57.1%, *SD* = 14.6%, chance 50%, *t*(102) = 4.95, *p* < .001). RAVLT delayed recall performance was *M* = 69.0% (*SD* = 19.2%).

SMT performance was significantly lower at visit 2 than at visit 1 (*t*(102) = 12.43, *p* < .001), demonstrating that memory for social information declined over the 21-day delay. To examine whether the decline in SMT performance across the 21-day delay differed by question subtype, we used a mixed effects ANOVA with visit and subtype as within-subjects factors and group as a between-subjects factor (Figure 3). Mauchly’s test indicated violations of sphericity for all effects involving subtype (all *p*’s < .001), so Greenhouse-Geisser corrected degrees of freedom are reported for those effects. There was a significant main effect of visit (*F*(1, 101) = 96.26, *p* < .001, η²= .063), reflecting the decline in performance across the delay, as well as a main effect of subtype (*F*(4.21, 424.85) = 14.51, *p* < .001, η²= .052). The visit × subtype interaction was significant (*F*(4.38, 442.87) = 12.31, *p* < .001, η²= .043), demonstrating that the magnitude of decline was not uniform across subtypes. Significant interactions were observed for group × visit × subtype (*F*(4.38, 442.87) = 8.17, *p* < .001, η²= .029) and group × subtype (*F*(4.21, 424.85) = 5.41, *p* < .001, η²= .020) but not group × visit (*F*(1, 101) = 0.04, *p* = .851, η² < .001). These effects suggest differences in subtype-level performance and decline across groups that may result from the different questions used at visits 1 and 2 for each group.

**Figure 3:**
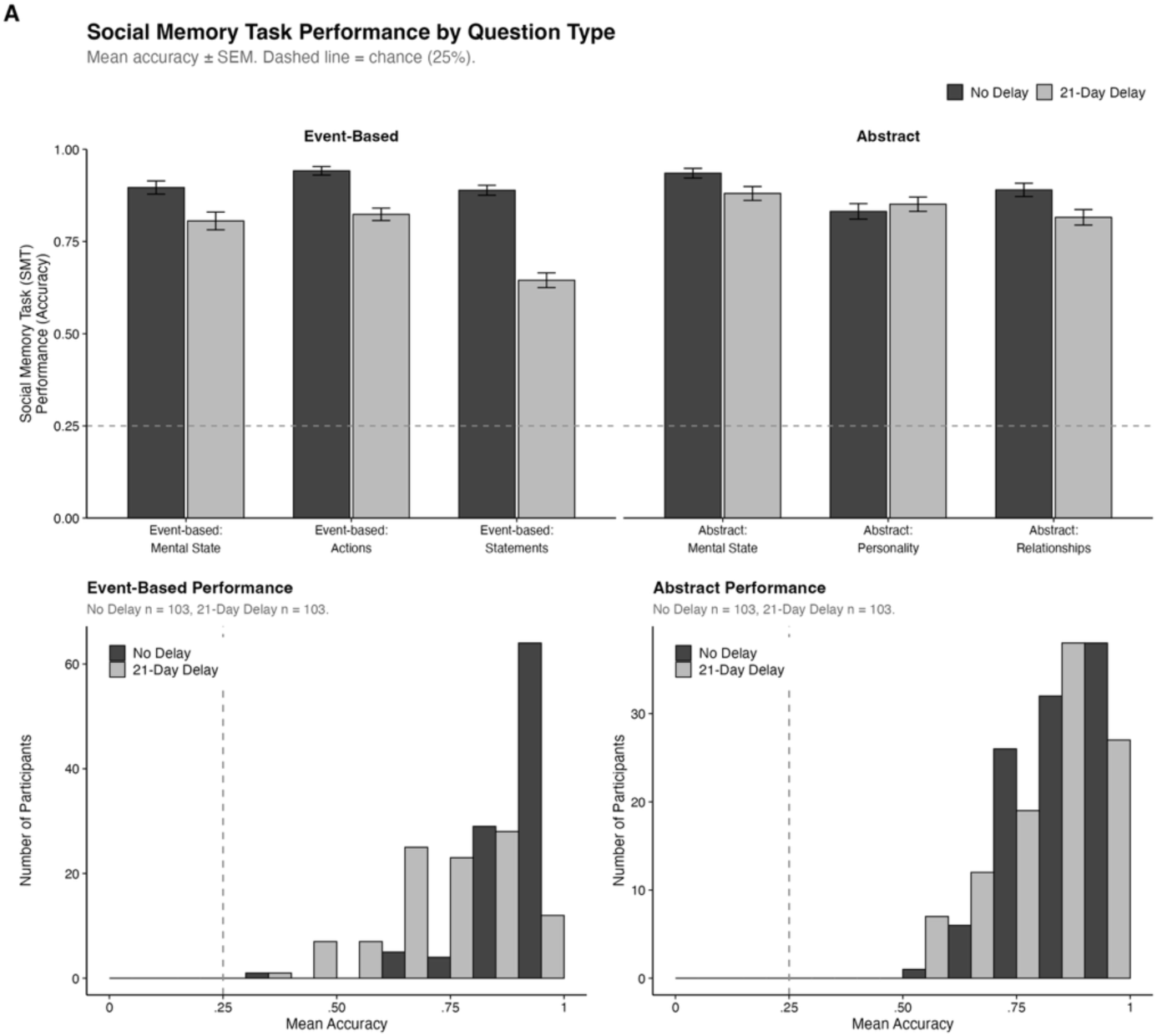
(A) Social memory task performance across visit 1 (no delay) and visit 2 (21-day delay) by question type. (B) Distributions of performance across participants for event-based questions. (C) Distributions for abstract questions.

To follow up the visit × subtype interaction, we conducted paired *t*-tests comparing visit 1 and visit 2 accuracy separately for each subtype. Among event-based subtypes, all three showed significant declines, including statements (*M*_Diff_ = 24.4%, *t*(102) = 10.88, *p* < .001), actions (*M*_Diff_ = 11.8%, *t*(102) = 6.54, *p* < .001), and mental states (*M*_Diff_ = 9%, *t*(102) = 3.59, *p* < .001). Among abstract subtypes, performance on mental state questions showed a decline (*M_Diff_* = 4.5%, *t*(102) = 2.34, *p* = .021), as did relationships (*M_Diff_* = 7.4%, *t*(102) = 2.63, *p* = .010), but not personality (*M_Diff_* = –1.9%, *t*(102) = −0.72, *p* = .476). These results suggest that memory for event-based information is susceptible to loss, whereas abstract information is largely retained across the delay.

### Internal Validity

We next measured split-half reliability to assess internal validity (Figure 4). To prevent variance attributable to group assignment from influencing reliability estimates, correlations were computed separately within each group, Fisher z-transformed, and averaged across groups. Signifiance was assessed with a permutation test: subject labels were permuted within each group, and correlations were computed as described above to build a null distribution (10,000 permutations). All split-half correlations were Spearman-Brown corrected.

**Figure 4:**
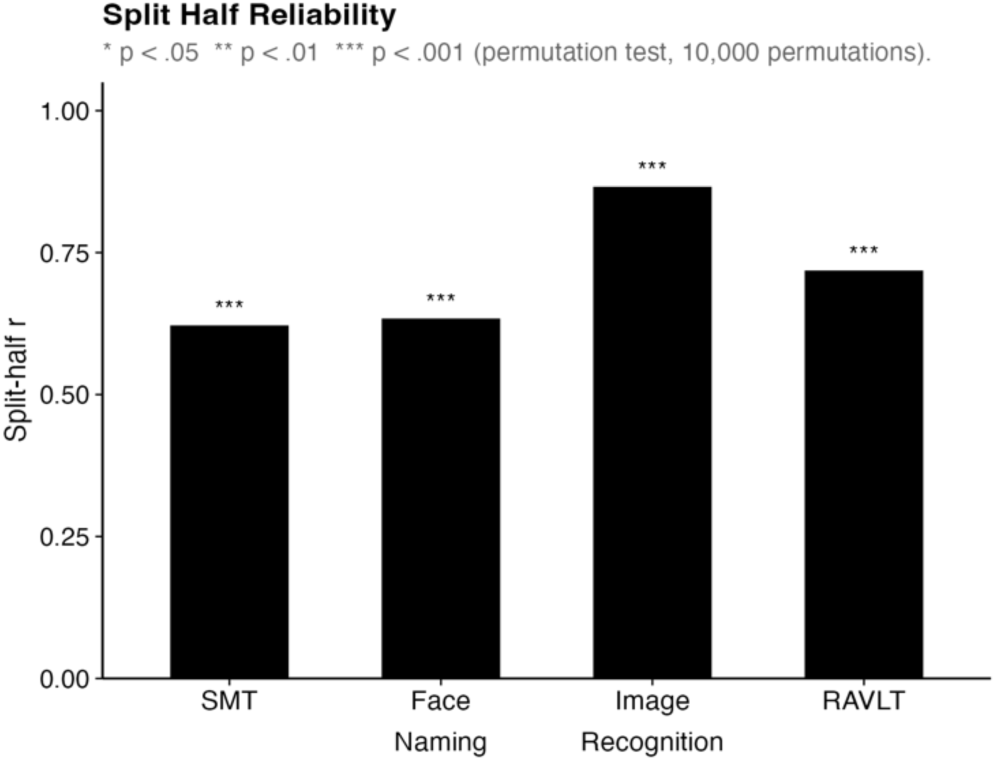
Split-half reliability (Spearman-Brown-corrected correlation) for the social memory task (SMT), image recognition, face naming, and Rey Auditory Verbal Learning Test (RAVLT) delayed recognition tasks.

SMT performance demonstrated moderate, significant internal reliability. This result was observed with data combined across visits (*r* = .62, *p* < .001), at visit 1 (*r =* .50, *p =* .002), and at visit 2 (*r =* .54, *p <* .001). The result was observed for event-related questions (r = .62, p < .001) but not abstract questions (r = .07, p = .726), indicating that reliable performance on the SMT derives largely from event-related questions. The remaining tasks also demonstrated moderate to high internal reliability: face naming (*r =* .63, *p <* .001), image recognition (*r =* .86, *p <* .001), and the RAVLT (*r =* .72, *p <* .001).

### Construct Validity

To examine the construct validity of the SMT, we measure correlations with the other memory measures (Figure 5, Supplemenary Figure 1, Supplementary Table 1). Significance was assessed using a permutation test (see Internal Validity). SMT visit 1 performance was correlated with face naming (*r =* .26, *p =* .007) and RAVLT (*r =* .22, *p =* .033), but not image recognition (*r =* .07, *p =* .514). SMT visit 2 performance was significantly correlated with performance on all three tasks (face naming: *r =* .39, *p <* .001; image recognition: *r =* .23, *p =* .023; RAVLT delayed recall: *r =* .20, *p =* .048). Critically, SMT was most strongly correlated across visits with the face naming task, suggesting a relationship between memory for high-level social information and face-name associations. Performance on the other three memory tasks was not correlated, including face naming and image recognition (*r =* .16, *p =* .109), face naming and RAVLT (*r =* .02, *p =* .835), and image recognition and RAVLT (*r =* .15, *p =* .130).

**Figure 5:**
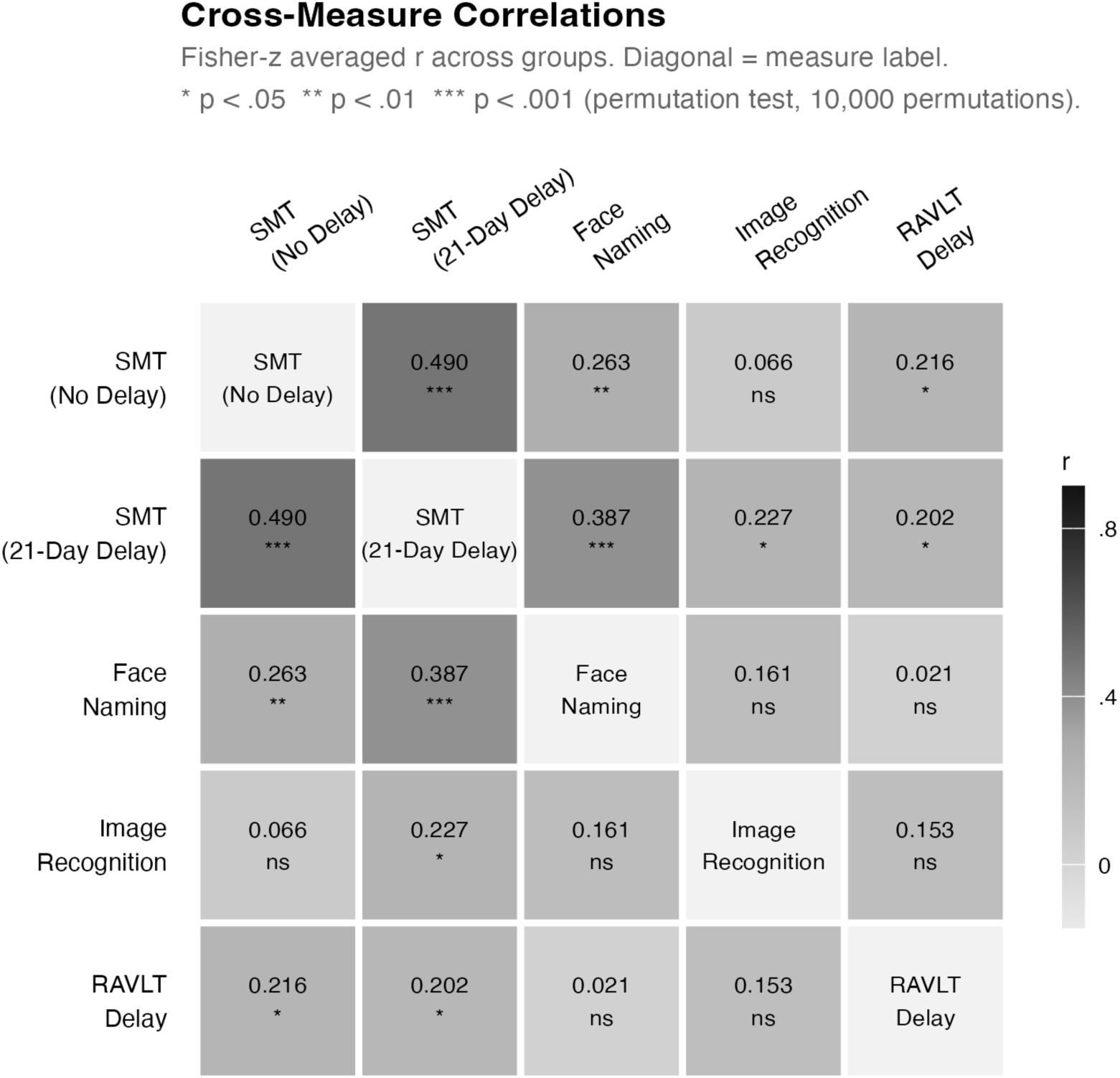
Correlation matrix across all tasks: the social memory task (SMT) visit 1 (no delay), visit 2 (21-day delay) image recognition, face naming, and Rey Auditory Verbal Learning Test (RAVLT) delayed recall.

Does performance across SMT question subtypes reflect a common underlying factor? We observed correlations between event-related question subtypes, but not abstract subtypes, or between event-related and abstract (detailed description in Supplementary Results, Supplementary Figure 2, Supplementary Table 2). These results suggest a common factor driving performance on event-related but not abstract questions.

Finally, we asked whether SMT captures variance across individuals that goes beyond what can be explained by the other memory measures. We computed the partial correlation between SMT performance at visits 1 and 2, controlling for face naming, image recognition, and RAVLT delayed recall. SMT performance was moderately and significantly correlated across visits (partial correlation *r* = .41, *p* < .001; full correlation *r* = .49, *p* < .001). This suggests that the individual differences measured by the SMT are not fully accounted for by established long-term memory measures.

## Discussion

The present study developed and evaluated a behavioral measure of long-term memory for naturalistic social content. Participants watched the pilot episode of *Friday Night Lights* and answered multiple choice questions probing memory for high-level social information, including characters’ actions, statements, mental states, personality traits, and relationships. The results provide a proof of concept for this approach to measuring individual differences in social memory, demonstrating moderate internal reliability, expected patterns of memory decline over time, convergent validity with established measures of episodic and associative memory, and divergence from visual recognition memory.

Performance on the SMT was well above chance (mean 84%, chance 25%), indicating that participants encoded social information from the video stimulus and that our multiple choice questions effectively probed this content. Performance declined over a 21-day delay from 90% to 78%, reflecting memory loss over time. Despite the decline, performance remained relatively high and well above chance after the delay, indicating that participants retained rich social content weeks after a single viewing of a naturalistic television episode. This is consistent with prior work demonstrating that memory for narrative video content is robust and long-lasting (Delarazan et al., 2023; Furman et al., 2007).

Larger performance declines were observed for event-related questions, about characters’ behavior and mental states tied to specific events in the narrative, than for abstract questions. This result is consistent with a large body of evidence showing that memory for details of events decays more quickly than memory for the abstracted gist of what happened (Brainerd & Reyna, 1990; Graesser et al., 1994; Loftus & Palmer, 1974; Schank & Abelson, 1977). Faster memory decay for actions (event-related) compared to personality traits (abstract) is consistent with work on spontaneous trait inference, arguing that personality traits are automatically inferred from others’ behavior and retained even as information about specific actions is lost (Carlston & Skowronski, 1994; Todorov & Uleman, 2003; Uleman et al., 1996). While prior work on impression formation has largely focused on personality traits inferred from actions, the current work indicates that similar dynamics are at play in memory for mental state information: larger decay was observed for event-related (9%) versus abstract (5.5%) questions about characters’ mental states. Though preliminary, these results indicate that initial detailed representations of characters’ mental states in the narrative (who felt what in which situation) give way over time to abstracted representations (who felt what in general).

A key criterion for a useful individual difference measure is that variation in performance reflects reliable differences across participants rather than noise. We found that the SMT had moderate, significant split-half reliability (*r* = .62). While slightly below the strong correlation range (*r* > .7) desired for a psychometric test, these results demonstrate that the SMT captures meaningful individual differences. Several results suggest modifications of the task that could improve reliability in future implementations. First, reliability in the present data may be diminished by ceiling effects, given high accuracies observed (especially for no delay performance; Fig. 3B-C). This issue could be resolved by increasing task difficulty or studying a population with a broader range of cognitive ability than the predominantly undergraduate young adult sample used here. Second, split-half reliability was substantially higher for event-related (*r* = .62) than abstract questions (*r* = .07). It remains unclear whether this reflects a genuine lack of reliable performance for abstract questions, or a larger ceiling effect, which could have resulted from the minimal decay observed for abstract questions. Regardless, this suggests that testing memory for social information tied to specific events may yield a more reliable individual difference measure than memory for abstract information.

To what extent is performance across SMT question subtype driven by a common underlying factor? Correlations across question subtypes were variable, with significant correlations observed within the event-related subtypes, but few significant correlations within abstract subtypes or between event-related and abstract. These results should be interpreted with caution, given 1) the small number of items included per subtype; 2) low reliability of abstract question performance; and 3) potential ceiling effects, as discussed above. Nevertheless, the results indicate that memory for event-related information about mental states, actions, and statements may be driven by a common factor. By contrast, performance on abstract questions was not sufficiently reliable to meaningfully assess convergence with other measures.

How does SMT performance relate to other long-term memory tasks, and what can this tell us about the cognitive processes that drive variation in the measure? Given that the SMT uses a complex stimulus (video narrative) and a complex behavioral measure (multiple choice questions), variation could in principle be driven by a variety of cognitive processes spanning the domains of vision, attention, social cognition, and memory. To constrain our interpretation of individual differences, we used a battery of additional long-term memory tasks, including two measures testing memory for other information from the same TV episode – face naming and image recognition – along with a standard, validated measure of verbal episodic memory, the RAVLT. SMT showed a structured pattern of correlations across tasks, revealing both convergence and divergence with other measures. The highest correlations were observed with face-name association memory (.26 – .39), along with weak but significant correlations with RAVLT (.20 – .22), and weak-to-no correlations with image recognition (.07 – .23). These results suggest that the SMT is related to distinct tests of relational long-term memory, while dissociable from visual recognition memory.

Correlations between the SMT and RAVLT delayed recall performance demonstrate convergent validity with an established measure of hippocampal-dependent long-term memory. These two tasks differ in content (word lists versus social narrative), modality (auditory/verbal versus audiovisual), and encoding context (intentional in the RAVLT versus incidental in the SMT). Shared variance can therefore not be explained by superficial task demands, and is better explained by a common underlying cognitive factor. By testing uncued recall that doesn’t rely on priming, recognition, or semantic association, RAVLT delayed recall provides a measure of episodic memory with variation that relates to hippocampal function (Aumont et al., 2025; Est⌈ vez−Gonz〈lez et al., 2003; Marchiani et al., 2008; Putcha et al., 2019). The SMT tests memory for relational information – relationships between characters and their mental states, traits, and behavior, as well as specific situations or contexts – that is also expected to be hippocampal-dependent (Cohen & Eichenbaum, 1993). Correlations were also observed between the SMT and face naming, another test of memory for relational social information, providing additional evidence for convergent validity with measures of hippocampal-dependent long-term memory (Rentz et al., 2011).

By contrast to its correlation with relational memory measures, the SMT did not show consistent correlations with image recognition. Likewise, face-name association memory did not show a significant correlation with image recognition, despite superficial similarities between the tasks: both involve responding to a visual image of a character’s face from *Friday Night Lights*. These dissociations may relate to differences in the cognitive and neural operations required to perform each task. While SMT and face naming both require memory for relations, image recognition only requires memory for a single object (the character’s face; Davachi, 2006; Ranganath & Ritchey, 2012). Visual recognition tasks can be solved using familiarity-based mechanisms thought to rely on perirhinal cortex rather than the hippocampus (Brown & Aggleton, 2001; Yonelinas, 2002). Furthermore, the image recognition task is purely visual, while both the SMT and face naming task involve verbal or conceptual information. Neural mechanisms supporting long-term memory for visual information about faces and conceptual information about familiar people are thought to be distinct, with the former reaching the hippocampus via perirhinal cortex, and the latter via association cortex (Deen & Freiwald, 2025). The observed dissociation between social memory and image recognition is therefore plausible given our understanding of the cognitive and neural mechanisms required for these tasks.

One limitation of the current study is its restricted sample demographics, with participants who are undergraduates at a selective university, within a narrow age range (18-22), and predominantly white. Future work should implement the SMT with participants with a broader range of demographics and cognitive ability. As discussed above, studying a more diverse population might reduce ceiling effects and increase reliability of the task. Using the SMT in aging or neurodegenerative populations could yield insight into how social memory is affected in cognitive decline. While *Friday Night Lights* is a high-school drama, we believe it would provide an engaging stimulus for a range of ages, given its compelling plot and focus on broad themes such as the pressure to succeed under scrutiny and the heartbreak of athletic injury.

A related limitation is that understanding the social content of *Friday Night Lights* may rely on cultural knowledge, such as understanding of the game of football, American high school football culture, or the role of sports in small-town community identity. In developing multiple choice questions, we tried to avoid questions that assumed specific prior knowledge, instead focusing on information that was clearly presented in the video narrative. Nevertheless, cultural knowledge may shape stimulus encoding and retention in ways that are difficult to measure and account for. We note that the methodological approach developed here – probing social content from a video narrative – could be used with other specific videos, to tailor our task to participants from another country and/or cultural background.

Overall, this study presents first steps toward the development of a psychometrically validated behavioral measure of social memory. We demonstrate the feasibility of using a narrative video stimulus combined with targeted multiple choice questions probing memory for social content. Despite the complexity of the task, we observed moderate reliability and a meaningful pattern of convergence and divergence with other tasks, lending support to the approach. Our results suggest refinements that can be made in future work to increase reliability, such as restricting the measure to questions about specific events in the narrative, and combatting ceiling effects. Given recent work arguing for DN-B as a social memory system in the human brain, future studies should relate SMT behavior to individual differences in DN-B anatomy and function. We hope that the current study provides the foundation for studying social memory as a distinct and important cognitive domain.

## Data & code availability

All data, stimulus materials, experimental code, and analysis code for this project are available at https://osf.io/kgy4n/overview.

## Author contributions

B.D. conceptualized the project and provided supervision. B.D. and D.R. developed the behavioral tasks. D.R. and R.K. collected data. T.M. and B.D. performed project administration, data curation, data analysis, and manuscript writing.

## Supplementary Information

### Supplementary Results

#### Performance on the social memory task across groups

At visit 1, group 1 performed at 91.2% (*SD* = 7.6%, *t*(55) = 65.48, *p* < .001, *d* = 8.75), and group 2 at 88.9% (*SD* = 8.7%, *t*(46) = 50.52, *p* < .001, *d* = 7.37). At visit 2, group 1 performed at 77.3% (*SD* = 9.7%, *t*(55) = 40.54, *p* < .001, *d* = 5.42), and group 2 at 79.0% (*SD* = 11.6%, *t*(46) = 31.82, *p* < .001, *d* = 4.64). The decline in performance across visits was significant in both group 1 (*M*s = 91.2% vs. 77.3%, *t*(55) = 10.48, *p* < .001, *d* = 1.40), and group 2 (*M*s = 88.9% vs. 79.0%, *t*(46) = 7.21, *p* < .001, *d* = 1.05).

#### Show ratings

Participants rated the show on average 7.65 out of 10 (*SD* = 1.39, range = 4-10, *n* = 90). Ratings were similar across groups (*M_Group1_* = 7.67, *SD* = 1.45, *n* = 48; *M_Group2_* = 7.64, *SD* = 1.33, *n* = 42). Show ratings were not collected for 13 of the 103 participants. These participants are excluded from our analysis of correlations with show ratings. To examine whether subjective enjoyment of the video stimulus related to task performance, we tested whether participants’ enjoyment ratings of the show were correlated with accuracy on each task measure. Show ratings were not significantly associated with overall SMT performance at either visit 1 (*r* = .11, *t*(88) = 1.06, *p* = .294) or visit 2 (*r* = .003, *t*(88) = 0.02, *p* = .981). Participants’ show ratings were also unrelated to face naming (*r* = −.08, *t*(88) = −0.74, *p* = .460), image recognition (*r* = .03, *t*(88) = 0.25, *p* = .801), and RAVLT Delay performance (*r* = −.04, *t*(88) = −0.37, *p* = .715). Taken together, these results suggest that performance on the SMT and related measures was not meaningfully influenced by how much participants enjoyed the show.

#### Question subtype correlations

We computed correlations for all 15 pairwise correlations among SMT subtypes averaged across visit 1 (no delay) and visit 2 (21-day delay). The strongest and most reliable association was between event-based: actions and event-based: statements (*r* = .36, *p* < .001). Five additional pairs were significantly correlated: event-based: mental states and event-based: actions (*r* = .21, *p* = .003), event-based: actions and abstract: relationships (*r* = .21, *p* = .004), event-based: mental states and event-based: statements (*r* = .21, *p* = .004), and event-based: statements and abstract: relationships (*r* = .25, *p* < .001), and event-based: statements and abstract: mental states also reached significance (*r* = .14, *p* = .046). The remaining nine pairs were not significantly correlated (all *p*’s ≥ .105), including all pairings among the three abstract subtypes (all |*r*|’s ≤ .12) and most event-related-to-abstract pairs.

**Supplementary Figure 1:**
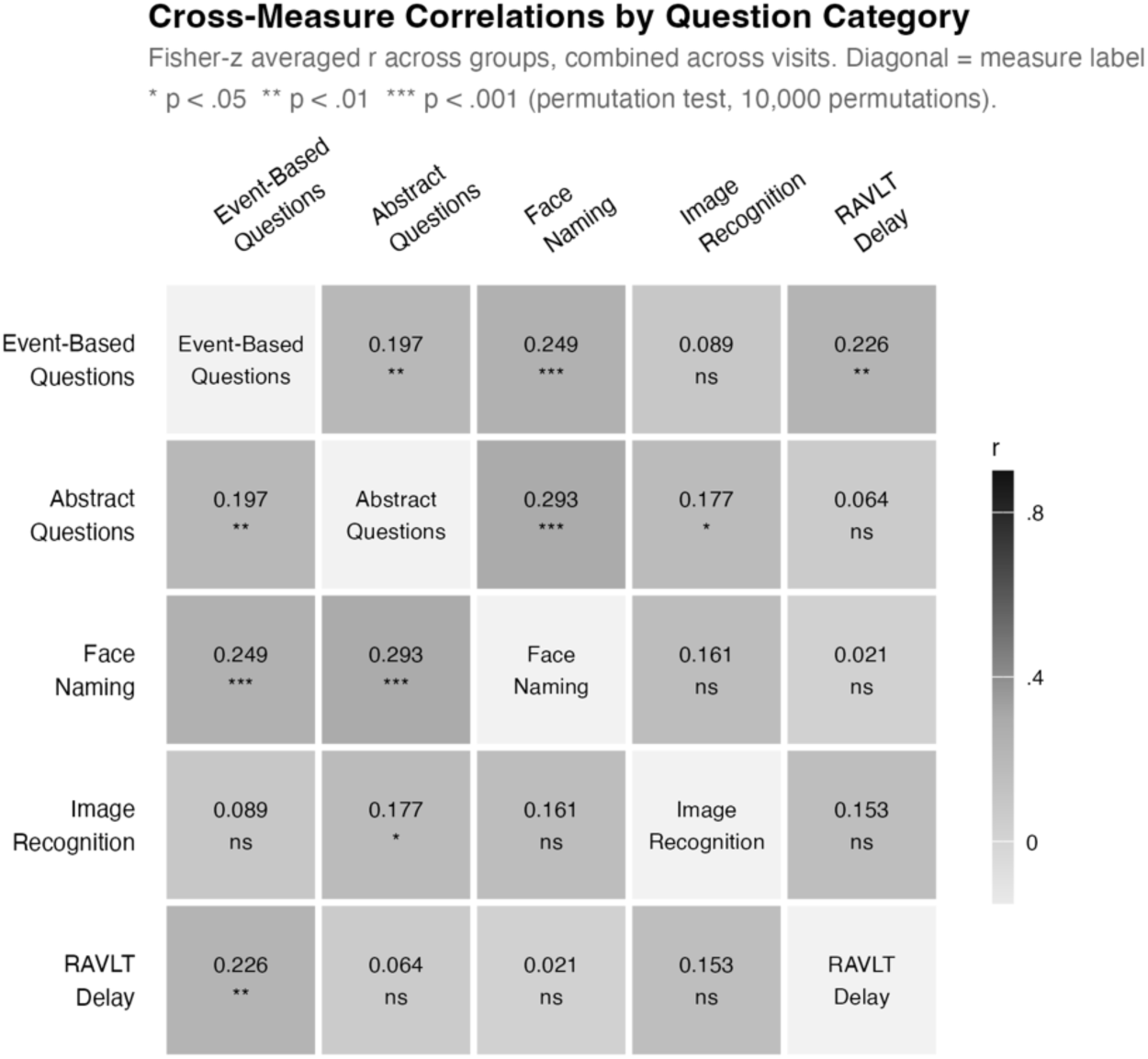
Correlation matrix across all tasks, with the social memory task separated by event-related and abstract questions.

**Supplementary Figure 2:**
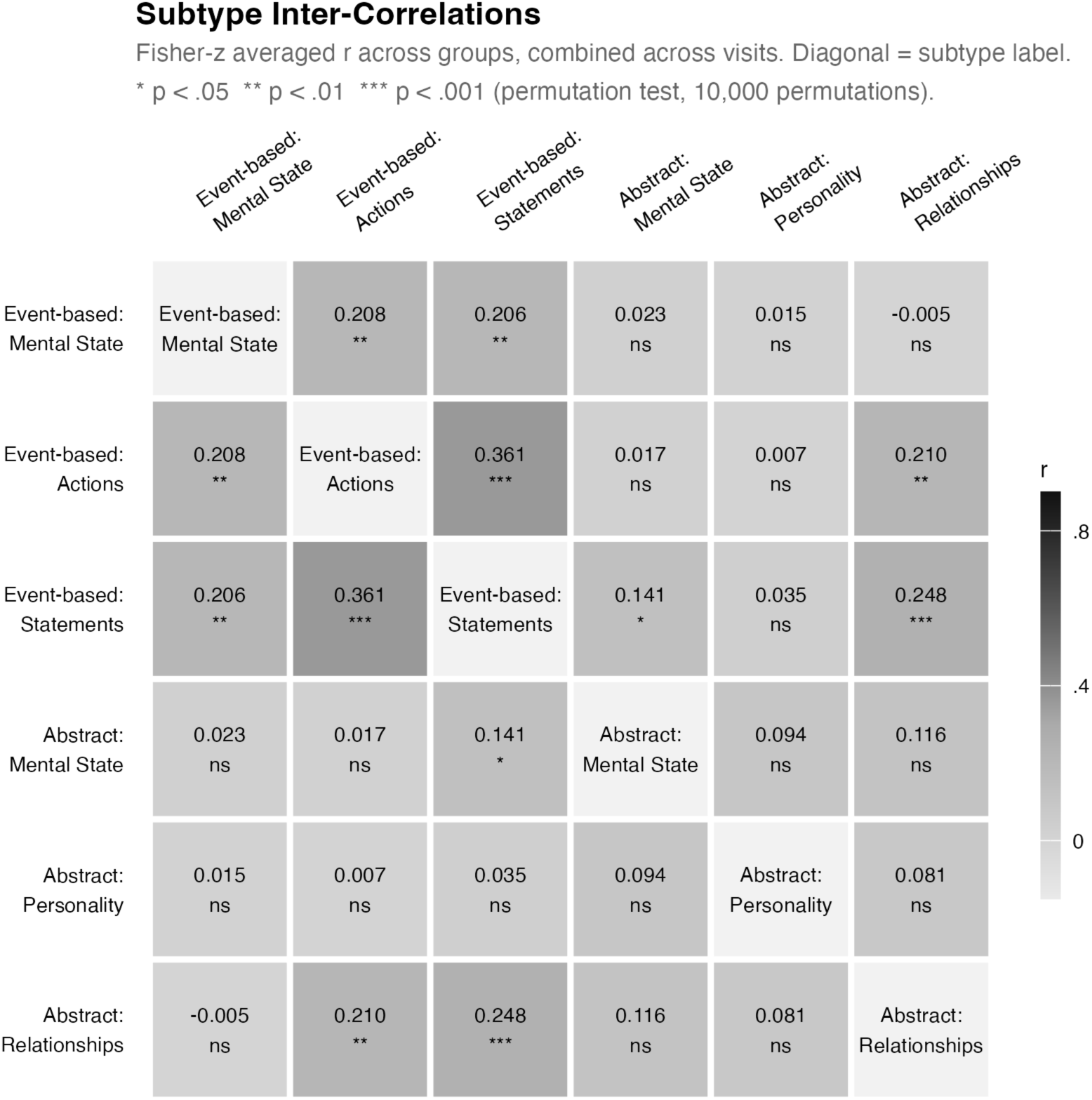
Correlation matrix across social memory task question subtypes.

**Supplementary Table 1:** Cross-task correlations, with social memory task separated by visit and question category.

| **Visit** | **Question Category** | **Measure** | **r** | **p** |
| --- | --- | --- | --- | --- |
| No Delay (V1) | Event | Face Naming | 0.179 | .072 |
| No Delay (V1) | Event | Image Recognition | 0.001 | .991 |
| No Delay (V1) | Event | RAVLT Delayed Recall | 0.285 | .003** |
| No Delay (V1) | Abstract | Face Naming | 0.239 | .018* |
| No Delay (V1) | Abstract | Image Recognition | 0.130 | .207 |
| No Delay (V1) | Abstract | RAVLT Delayed Recall | -0.048 | .638 |
| 21-Day Delay (V2) | Event | Face Naming | 0.316 | .002** |
| 21-Day Delay (V2) | Event | Image Recognition | 0.175 | .085 |
| 21-Day Delay (V2) | Event | RAVLT Delayed Recall | 0.164 | .101 |
| 21-Day Delay (V2) | Abstract | Face Naming | 0.346 | < .001*** |
| 21-Day Delay (V2) | Abstract | Image Recognition | 0.224 | .024* |
| 21-Day Delay (V2) | Abstract | RAVLT Delayed Recall | 0.174 | .082 |

Stars: \* $p < .05$ , \*\* $p < .01$ , \*\*\* $p < .001$

**Supplementary Table 2:** Social memory task subtype correlations, separated by visit.

| Visit | Subtype A | Subtype B | r | p |
| --- | --- | --- | --- | --- |
| No Delay (V1) | Event: Mental State | Event: Actions | 0.325 | .002** |
| No Delay (V1) | Event: Mental State | Event: Statements | 0.309 | .002** |
| No Delay (V1) | Event: Mental State | Abstract: Mental State | -0.080 | .415 |
| No Delay (V1) | Event: Mental State | Abstract: Personality | 0.109 | .281 |
| No Delay (V1) | Event: Mental State | Abstract: Relationships | 0.056 | .575 |
| No Delay (V1) | Event: Actions | Event: Statements | 0.353 | < .001*** |
| No Delay (V1) | Event: Actions | Abstract: Mental State | -0.008 | .913 |
| No Delay (V1) | Event: Actions | Abstract: Personality | -0.027 | .776 |
| No Delay (V1) | Event: Actions | Abstract: Relationships | 0.143 | .153 |
| No Delay (V1) | Event: Statements | Abstract: Mental State | 0.044 | .699 |
| No Delay (V1) | Event: Statements | Abstract: Personality | 0.073 | .483 |
| No Delay (V1) | Event: Statements | Abstract: Relationships | 0.185 | .066 |
| No Delay (V1) | Abstract: Mental State | Abstract: Personality | 0.108 | .260 |
| No Delay (V1) | Abstract: Mental State | Abstract: Relationships | 0.134 | .202 |
| No Delay (V1) | Abstract: Personality | Abstract: Relationships | -0.022 | .801 |
| 21-Day Delay (V2) | Event: Mental State | Event: Actions | 0.085 | .411 |
| 21-Day Delay (V2) | Event: Mental State | Event: Statements | 0.098 | .341 |
| 21-Day Delay (V2) | Event: Mental State | Abstract: Mental State | 0.126 | .211 |
| 21-Day Delay (V2) | Event: Mental State | Abstract: Personality | -0.080 | .438 |
| 21-Day Delay (V2) | Event: Mental State | Abstract: Relationships | -0.066 | .522 |
| 21-Day Delay (V2) | Event: Actions | Event: Statements | 0.369 | < .001*** |
| 21-Day Delay (V2) | Event: Actions | Abstract: Mental State | 0.042 | .681 |
| 21-Day Delay (V2) | Event: Actions | Abstract: Personality | 0.040 | .707 |
| 21-Day Delay (V2) | Event: Actions | Abstract: Relationships | 0.276 | .007** |
| 21-Day Delay (V2) | Event: Statements | Abstract: Mental State | 0.235 | .019* |
| 21-Day Delay (V2) | Event: Statements | Abstract: Personality | -0.003 | .975 |
| 21-Day Delay (V2) | Event: Statements | Abstract: Relationships | 0.310 | < .001*** |
| 21-Day Delay (V2) | Abstract: Mental State | Abstract: Personality | 0.080 | .425 |
| 21-Day Delay (V2) | Abstract: Mental State | Abstract: Relationships | 0.099 | .335 |
| 21-Day Delay (V2) | Abstract: Personality | Abstract: Relationships | 0.182 | .064 |

Stars: \* $p < .05$ , \*\* $p < .01$ , \*\*\* $p < .001$

